# PhaeoEpiView: An epigenome browser of the newly assembled genome of the model diatom *Phaeodactylum tricornutum*

**DOI:** 10.1101/2022.07.29.502047

**Authors:** Yue Wu, Chaumier Timothée, Eric Manirakiza, Alaguraj Veluchamy, Leila Tirichine

## Abstract

**Motivation:** Recent advances in DNA sequencing technologies in particular of long reads type greatly improved genomes assembly leading to discrepancies between both published annotations and epigenome tracks which did not keep pace with new assemblies. This comprises the availability of accurate resources which penalizes the progress in research.

**Results:** Here, we used the latest improved telomere to telomere assembly of the model pennate diatom *Phaeodactylum tricornutum* to lift over the gene models from Phatr3, a previously annotated reference genome. We used the lifted genome annotation including genes and transposable elements to map the epigenome landscape, namely DNA methylation and post translational modifications of histones providing the community with PhaeoEpiView, a browser that allows the visualization of epigenome data as well as transcripts on an updated reference genome to better understand the biological significance of the mapped data on contiguous genome rather than a fragmented one. We updated previously published histone marks with a more accurate mapping using monoclonal antibodies instead of polyclonal and deeper sequencing. PhaeoEpiView will be continuously updated with the newly published epigenomic data making it the largest and richest epigenome browser of any stramenopile. We expect that PhaeoEpiView will be a standard tool for the coming era of molecular environmental studies where epigenetics holds a place of choice.

**Availability:** PhaeoEpiView is available at: https://PhaeoEpiView.univ-nantes.fr

## Introduction

The genome of the model diatom *Phaeodactylum tricornutum* CCAP 1055/1 and the corresponding annotation were published in 2008 using whole genome shotgun paired-end Sanger sequencing (NCBI assembly ASM15095v2) (Bowler et al., 2008). Subsequently, Phatr3 annotation updated gene repertoire to introduce more than thousand novel genes and performed a comprehensive de novo annotation of repetitive elements showing novel classes of transposable elements using 90 RNA-Seq datasets combined with published expressed sequence tags and protein sequences (Rastogi et al., 2018). The first assembly of the genome contained 33 scaffolds among which 12 telomere-to-telomere chromosomes. Using long read sequencing, Filloramo et al., re-examined *P. tricornutum* assembly which led to additional sequence information but did not improve the continuity and chromosome-level scaffolds compared to the original reference genome (Filloramo et al., 2021). Recently, an approach combining long reads from the Oxford Nanopore minION platform and short high accurate reads from the Illumina NextSeq platform was used to perform a new assembly of *P. tricornutum* genome which led to 25 nuclear chromosomes improving thus the assembly (Giguere, 2021). However, Phatr3 annotation of *P. tricornutum* was not revisited in light of the new 25 telomeric chromosomes assembly which is often observed for most species where the annotations do not keep pace with new/improved assemblies.

*P. tricornutum* is an established model diatom widely used by an increasing community for fundamental research and biotech applications. Diatoms are one of the most abundant and highly diverse mostly photosynthetic eukaryotes, contributing to 20-25% of the Earth’s global carbon dioxide fixation (Field et al., 1998) and their photosynthetic activity accounts for about 40% of the marine primary production (Armbrust, 2009; Falkowski et al., 1998). Diatoms are highly successful and widely spread occupying large territories including marine, freshwater, sea ice, snow and even moist terrestrial habitats.

While whole-genome sequencing is critical to better understanding the ecological success of diatoms, primary sequence is only the basis for understanding how to read genetic programs, another layer of heritable information superimposed on the DNA sequence is epigenetic information. It has already been proposed that the ecological success of phytoplankton is also due to the adaptive dynamics conferred by epigenetic regulation mechanisms because point mutation-based processes may be too slow to permit adaptation to a dynamic ocean environment (Tirichine and Bowler, 2011). The epigenetic changes may lead to chromatin modifications, which may cause a stable alteration in transcriptional activity even after withdrawal of the triggering stress (Avramova, 2015). Pioneering work drew a comprehensive map of epigenetic marks including several permissive and repressive PTMs and DNA methylation in *P. tricornutum* and showed their contribution to mediate the response of diatom cells to environmental factors (Rastogi et al., 2015; Veluchamy et al., 2013; Veluchamy et al., 2015).

An important molecular tool box is available in *P. tricornutum* including epigenomic data which are only found in the partial assembly. To make such a resource available on the newly assembled genome, we used the new 25 to 25 telomere assembly to map the epigenetic data including Post-translational modifications of histones (PTMs) and DNA methylation that were previously published (Hoguin, 2021; Veluchamy *et al*., 2013; Veluchamy *et al*., 2015). Prior to this, we lifted the Phatr3 annotation using a gene based approach.

### Implementation

PhaeoEpiView was built using two steps (i) Phatr3 gene annotation lifting onto the new 25 chromosomes assembly (Phatr4) (ii) mapping of the previously published epigenetic marks and transcripts on the new assembly. For more accuracy and homogeneity of the used data, chromatin immunoprecipitation with deep sequencing was carried out using monoclonal antibodies to replace two marks, H3K9me3 and H3K27me3 for which polyclonal antibodies were used in the previous study.

In the first step, instead of whole genome-based comparison, we adapted a gene-based sequence alignment for lifting the annotation from Phatr3 to Phatr4 assembly. Features such as mRNA, CDS and exons from the reference Phatr3 were used to infer genes and transcripts in target assembly. Using minimap2 and Liftoff tools, exons are aligned first to preserve the gene structure of the Phatr3 annotation (Li, 2018; Shumate, 2021). Minimap is used with 50 secondary mappings, end bonus of 5 and chaining score of 0.5. Genes are lifted and considered mapped successfully if the alignment coverage and sequence identity in the child features (usually exons/CDS) is >= 50%. Unplaced genes and genes with extra copy number are tagged and separated (Supplementary Table 1). Out of the 12178 genes from Phatr3 annotation, 11739 were lifted successfully (Supplementary File 2).

In order to validate the lifted annotation, we aligned RNAseq reads to both the previous and the new genome version then compared every gene quantification. The vast majority of them had a difference of quantification (Supplementary Figure 1) and length lower than +/-0.1% between Phatr3 and Phatr3_lift (Phatr4). Missing genes were then examined: 178 out of 439 (40%) were found to be located on short regions that are no longer present in Phatr4 assembly according to whole-genome alignment provided in (Giguere, 2021), half of them clustered on previous chr_5 and chr_21 (Supplementary Table 1). Most of the remaining 261 missing genes showed similarity to already lifted genes suggesting that they are either duplicated or allelic.

In the second step, transposable elements annotation available from (Giguere, 2021) was added to PhaeoEpiView as Phatr4 TEs track. Finally, previously published expression data at two different time points, DNA methylation and PTMs tracks were implemented in the browser and systematic comparison was made with the previous assembly mapping using unchanged regions as anchors (Supplementary File 1). PhaeoEpiView was implemented as a Jbrowse2 instance (Buels et al., 2016) and made public on a virtual machine hosted at Nantes University datacenter. It can currently display one track for each of the genes, TEs, transcript levels, McrBC and Bisulfite-seq DNA methylation and five histone PTMs (H3K9/14Ac, H3K4me2, H3K4me3, H3K9me2, H3K9me3, H3K27me3). The browser will be regularly updated with relevant epigenomic data when published in the future, making PhaeoEpiView a live platform for a comprehensive genomic and epigenomic resource of the model microalgae *P. tricornutum*.

## Conclusion

PhaeoEpiView is an open source browser that provides an up to date genome and epigenome view of the model diatom *Phaeodactylum tricornutum*. With the lifted genes annotation, the epigenome and transcriptome landscapes can be visualized on a fully assembled genome providing an accurate view of epigenetic regulation of genes and TEs which was incomplete on the previously fragmented genome. PhaeoEpiView allows users to upload their own tracks in private session for visualization and data interpretation purposes. PhaeoEpiView is intuitive, easy to use and represent the first epigenome browser of a photosynthetic unicellular species which will undoubtedly contribute to boost research in microalgae and single celled species in general.

**Figure 1.**
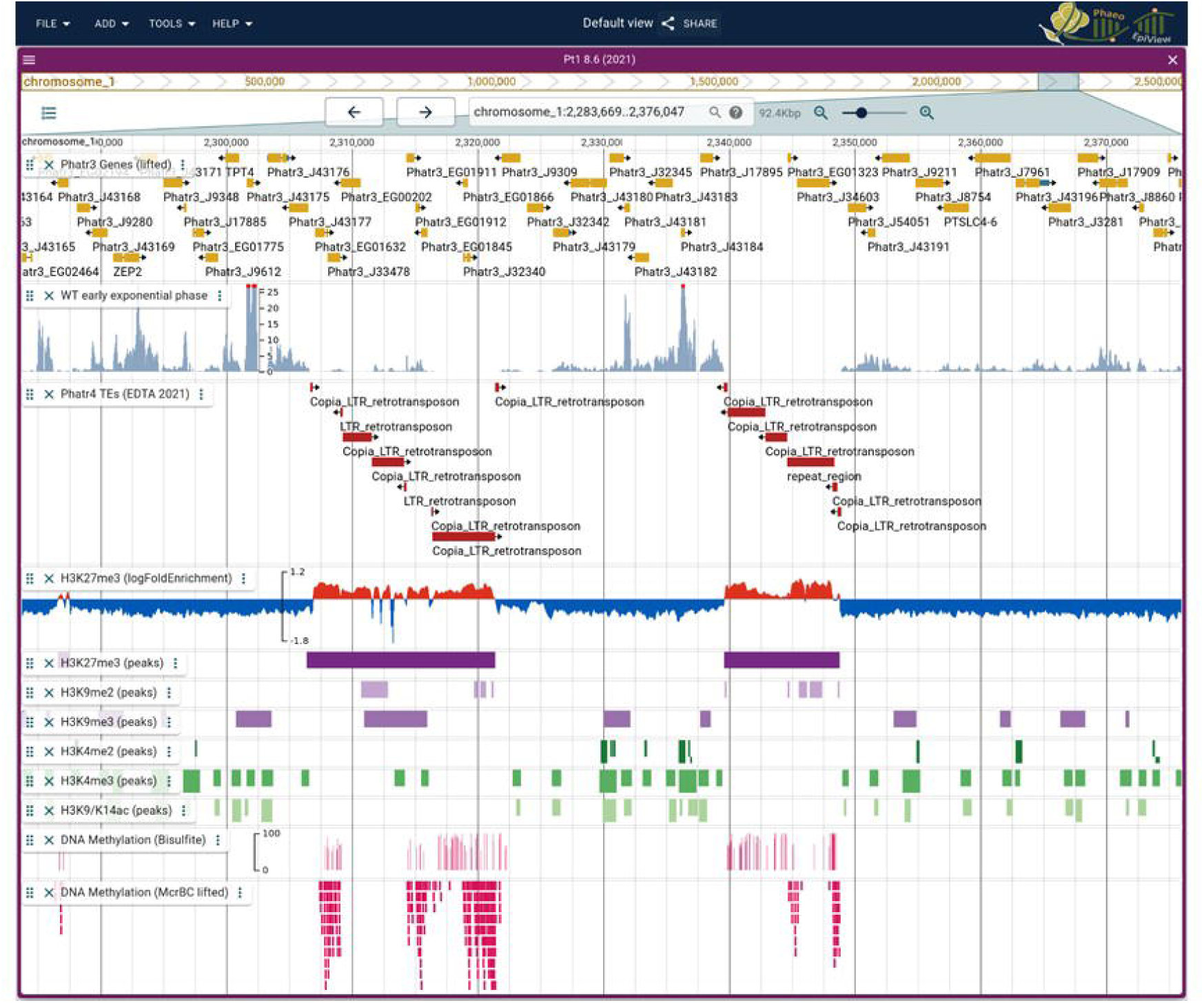
Snapshot of PhaeoEpiView browser illustrating the different tracks of genes, transposable elements, histone marks and DNA methylation. Both peaks and log2 fold enrichment between IP and Input are displayed for H3K27me3.

## Acknowledgements

This work was supported by the Epicycle ANR project (ANR-19-CE20-0028-02) and Région Pays de la Loire (ConnecTalent EPIALG project) to LT. YW was supported by a PhD fellowship from the Chinese Scholarship Council (CSC-201908340073). We are grateful to the Bioinformatics Core Facility of Nantes University (BiRD-Biogenouest) for its technical support.

## Supplementary methods

### Culture and growth conditions

*Phaeodactylum tricornutum* Bohlin Clone Pt1 8.6 (CCMP2561) cells were obtained from the culture collection of the Provasoli-Guillard National Center for Culture of Marine Phytoplankton (Bigelow Laboratory for Ocean Sciences, USA). Constantly shaken (100 rpm) cultures were grown at 19°C, 60 µmol photons m^-2^ s^-1^ and with a 12h light / 12h dark photoperiod in sterile Enhanced Artificial Sea Water (EASW) medium (Vartanian, et al., 2009). For Chromatin immunoprecipitation-sequencing, cultures were seeded at 50.000 cells/ml in duplicate and grown side by side in 1000 ml erlens until early-exponential at 10^6^ cells/ml. Culture growth was measured using a hematocytometer (Fisher Scientific, Pittsburgh, PA, USA).

### Chromatin extraction and immunoprecipitation

Chromatin isolation was performed as described previously (Lin, et al., 2012) with few modifications. Briefly, the incubation step in buffer II is repeated several times until the pellet becomes white. Each ChIP-seq experiment was conducted in two independent biological replicates. Monoclonal antibodies from Cell Signaling Technology were used for immunoprecipitation, H3K9me3 (13969), and H3K27me3 (9733).

### ChIP-Seq analysis

Pair-end sequencing of H3K9me3 and H3K27me3 ChIP and input samples was performed on Illumina NovaSeq platform with read length of 2 × 150 bp. Previously published ChIP sequencing for H3K9me2, H3K9me3, H3K4me2, H3K27me3, H3K9/K14Ac and H3K4me3 were retrieved from NCBI’s Gene Expression Omnibus accessions GSE68513 and GSE139676 (Veluchamy, et al., 2015; Zhao, et al., 2021). Raw reads were filtered and low-quality read pairs were discarded using Trim Galore 0.6.7 (https://doi.org/10.5281/zenodo.5127899) with a read quality (Phred score) cutoff of 20 and a stringency value of 3bp. Using the 25 to 25 telomere assembly published in 2021 as a reference genome, the filtered reads were mapped using Bowtie2 2.4.5 (Langmead and Salzberg, 2012). We then performed the processing and filtering of the alignments using Samtools 1.15 “fixmate -m” and “markdup -r” modules (Danecek, et al., 2021). Two biological replicates for each ChIP were performed and read counts showed a good Pearson correlation by Deeptools multiBamSummary v3.5.1 with a bin size of 1000bp (Ramirez, et al., 2014). To identify regions that were significantly enriched, we used MACS2 v2.2.7.1 (Zhang, et al., 2008) on the combination of the two replicates with “callpeak –qvalue 0.05 --nomodel --SPMR --bdg” options. In addition, extension size was set to the arithmetic mean of the two IP replicates fragment size for each mark, as determined by MACS2 predicted module with “-m 2 70” MFOLD value. Furthermore, “--broad” calling mode was activated for H3K9me2 and H3K9me3 that were previously described as broad histone marks. For the narrow marks H3K4me2, H3K9/K14Ac and H3K4me3, peaks summits were called with “-- call-summits”. Following previously published work, SICER2 v1.0.3 (Zang, et al., 2009) was used with “-w 200 -g 600 -fdr 0.05” to call peaks for H3K27me3.

Output normalized Fold Enrichment signal files were generated with MACS2 “bdgcmp” module and transformed to BigWig using Deeptools bedGraphToBigWig. Then, Pearson correlation between our new data and previously published data for H3K9me3 and H3K27me3 was performed using Deeptools plotCorrelation.

### Expression analysis

Early and late exponential growth phase Illumina RNA-seq data (SRR5274697, SRR5274696, SRR5274695 and SRR5274694) from (Murik, et al., 2019) were trimmed using Trim Galore 0.6.7 with a read quality (Phred score) cutoff of 20 and a stringency value of 3bp. Technical replicates were merged and mapped to the reference assembly with STAR 2.7.10a (https://www.ncbi.nlm.nih.gov/pubmed/23104886). Primary alignments only were processed with Deeptools bamCoverage 3.5.0 with “--normalizeUsing BPM --ignoreDuplicates -- centerReads” to generate normalized coverage files to be displayed in PhaeoEpiView.

### DNA methylation analysis

McrBC DNA methylation annotation data from (https://www.nature.com/articles/ncomms3091) was lifted from the previous assembly to the new 25 chromosomes using Liftoff. Bisulfite sequencing data (Hoguin, 2021) were processed with Bismark v0.22.3 (https://pubmed.ncbi.nlm.nih.gov/21493656/) and methylated regions having less than 50% methylated reads or less than 5 supporting reads were filtered out.

## Legend

**Supplementary Figure 1**. Comparison of RNA-seq quantification per gene on the 2008 (33 scaffold/chromosomes) and 2021 (25 chromosomes) assembled genomes. Two RNA-seq replicates were used.

**Supplementary Table 1**. List of the genes not recovered on the lifted annotation with their coordinates and features.

**Supplementary File 1**. Supplementary materials and methods.

**Supplementary File 2**. GFF3 file annotating *Phaeodactylum tricornutum* 2021 assembly with Phatr3 lifted genes.

