## Supplementary figures and images for "PhaeoEpiView: An epigenome browser of the newly assembled genome of the model diatom *Phaeodactylum tricornutum*"

### Supplementary figure 1

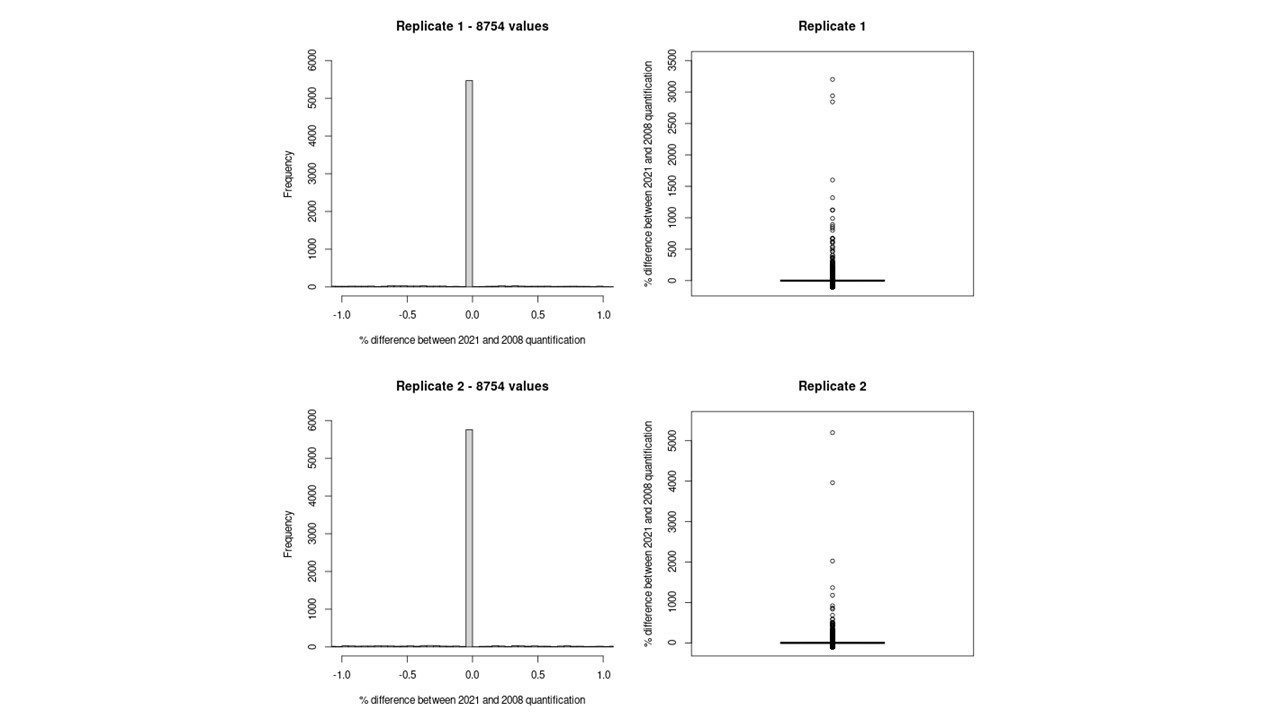
